# Co-evolutionary patterns shown in *Ostreococcus*-virus system from the Western Baltic Sea in freshly isolated hosts and viruses

**DOI:** 10.1101/2023.01.30.526186

**Authors:** Luisa Listmann, Carina Peters, Janina Rahlff, Sarah P. Esser, Elisa Schaum

## Abstract

Marine viruses are a major driver of phytoplankton mortality and thereby influence biogeochemical cycling of carbon and other nutrients. In recent years, an understanding of the potential importance of phytoplankton-targeting viruses on ecosystem dynamics has emerged, but experimental investigations of host-virus interactions on a broad spatial and temporal scale are still missing. Here, we investigated in detail a phytoplankton host’s responses reacting to infections by species-specific viruses from i) distinct geographical regions and ii) different sampling seasons. Specifically, we used two species of picophytoplankton (1 µm) *Ostreococcus tauri* and *O. mediterraneus* and their viruses (size ca. 100 nm), which represent systems well-known in marine biology, but almost entirely ignored in evolutionary biology. The strains stem from different regions of the Southwestern Baltic Sea that vary in salinity and temperature. Using an experimental cross-infection set-up, we show that in this natural system evolutionary history, and thus the timing of when hosts and their associated viruses coexisted, was the main driver of infection patterns. In addition species and strain specificity underline the present understanding of rapid host-virus co-evolution.

## Introduction

Viruses and their interactions with hosts are omnipresent in the marine environment. In phytoplankton communities viruses exert strong top-down control. Specifically, within the microbial loop [1], viruses can infect both hetero- and autotrophic microbes [2]. ”shunting” carbon away from the classical food web. Consequently, viruses can have far reaching effects on biogeochemical cycles [3]. To date, the biology and diversity of phytoplankton viruses is mainly studied *via* cultivation, independent genomic analyses or visualization methods like virusFISH (fluorescence in situ hybridization), often on the single-virus or single-cell level [4, 5]. This is facilitated because prediction and binning tools (genomic analyses) for eukaryotic viruses have significantly advanced [6, 7]. Only in few cases, infections of viruses are investigated by cultivation approaches [8, 9]. Viruses are released into the water through host cell lysis, which explains why the presence of viruses is tied to the presence of hosts [10]. However, we still know very little about the underlying ecology of viral infections in phytoplankton.

Studying the ecology of viruses comprises both biotic and abiotic aspects. Specifically, we need to understand how strong and specific infections are or how often and under what conditions they happen. Different conditions could originate from seasonal changes that affect the temperature, nutrient environment, and host abundance and diversity. Studies about the biotic aspects of viral ecology have recently been conducted in a few phytoplankton species. Similar to bacteria-bacteriophage interactions, infections are highly species or even strain specific [8, 11, 12] and diversity within species could affect the host range of infections. Effects of abiotic factors have to date not been well studied but are of great importance since environmental conditions in particular have changed in the past and are going to change drastically in the future [13–15].

In this study, we focus on a pico-phytoplankton species, namely *Ostreococcus* sp., which has a size of about 1 µm and thus is the smallest known free-living eukaryote [16]. The *Ostreococcus* sp. species complex is a feasible and ecologically relevant model organism for laboratory studies due to its global distribution and prevalence in many marine environments [17]. *Ostreococcus* are infected by large prasinoviruses (family Phycodnaviridae). These double-stranded DNA viruses are 100 nm in size, and belong to the Nucleocytoplasmic Large DNA Viruses (NCLDV) family. Like their hosts, they have been found globally [18]. The infection style within the *Ostreococcus*-virus system is lytic with the potential for rapid development of resistance in the host and has been described in several recent studies that also investigated genetic mechanisms of the dynamics [11, 19]. However, these studies have mainly focused on a confined geographical region in the Mediterranean Sea and/or time of year and did not investigate how host-virus interactions can change in terms of infectability and resistance patterns within few (i.e. months) generations.

Here, we further investigated the biology of the *Ostreococcus*-virus system by studying the infection patterns of both hosts and viruses that have experienced different evolutionary histories. The Baltic Sea lends itself well for investigating environmental gradients within geographical regions that are close enough to mitigate confounding effects, such as light and nutrient availability. This brackish sea is characterized by North-East to South-West gradients of both salinity and temperature as well as variability therein [20]. In the future, the Baltic Sea is predicted to become both sweeter and warmer, meaning that comparing different conditions present today will already allow for some insights into what we may expect in the future [21]. Within the *Ostreococcus*-virus system from the Baltic Sea, we are particularly interested in the following questions: i) when and where are *Ostreococcus* and its viruses likely present? ii) How species or strain specific are infections in the Baltic Sea *Ostreococcus*-virus system? and iii) How do infection patterns change over time or between geographical regions?

## Material and Methods

### Ostreococcus and virus isolations

In addition to previously isolated and studied strains of *Ostreococcus* [22], we isolated further 20 novel strains of *Ostreococcus mediterraneus* and 80 new viral strains. During seven cruises from 2018 to 2020 (AL505, AL513, AL520, AL521, AL522, AL524, AL530 and Assemble Plus project MIDSEAS), we collected water samples at 105 stations (Figure 1) in the Western Baltic Sea. The regions of the Baltic Sea included 4 different areas: the coastal Swedish Skagerrak (S) close to Tjärnö Marine Station [23], the Kiel area (KB and MB, Kiel Bight and Mecklenburg Basin, respectively), the Arkona Basin (AK) and Bornholm Basin (BB).

**Figure 1.**
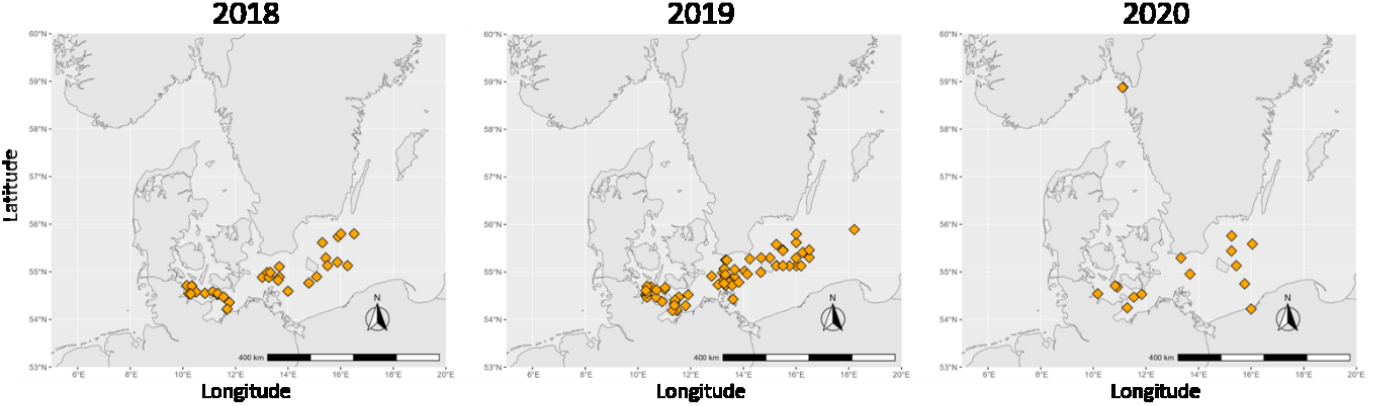
The three panels show the stations in the Western Baltic Sea that were sampled during three consecutive years. The sampling effort in 2019 was the highest with five cruises in total.

The water samples were taken at depths between 1 and 5 meters. The water was immediately filtered through a 35 µm mesh (Hydrobios ®) to remove grazers and further 2 L of two size fractions each were collected: first 2 L of filtrate through 0.45 µm (Whatman ®) were taken for later processing in the home laboratory in Hamburg, Germany. Second, 2 L of 2 µm filtrate were concentrated to 200 mL, of which 40 mL were taken for later processing in the home laboratory. The 0.45 µm filtrate was concentrated using tangential filtration (100’000 MWCO PES Sartorius ®) and analysed *via* flow cytometry to detect viral populations in the sample (see SI for flow cytometry description).

*Ostreococcus* spp. were isolated using the isolation by dilution method [24]. Upon successful isolation of *Ostreococcus* strains, viruses were isolated the following way: 20 mL of each *Ostreococcus* strain was combined with 1 mL of the 0.45 µm concentrated water sample of the respective geographical area (see Fig. S3; all combinations of hosts and virus water samples in upper panel, grey boxes) resulting in ca. 250 combinations. After 5, 10 and 14 days, the cell numbers were counted via flow cytometry (see below) and any *Ostreococcus* culture that had ≤ 50% cells than a control, was considered lysed (Fig. S3, green boxes in upper panel). These cultures were then filtered through a 0.45 µm filter membrane (Whatman ®), and the lysate was used for a second round of liquid infection. The lysis was checked again as before, and after a second successful infection, the lysate was used to isolate single virus strains using a plaque assay [25]. Two rounds of plaque formation and picking followed in semi-solid 0.2% agarose (Thermofisher ®) after which single viral strains were obtained (Fig. S3, remaining plaque assays in lower panel).

### Species identity and diversity

For DNA extraction, 2 mL of exponentially growing *Ostreococcus* culture was pelleted via centrifugation at 8’000 g. The pellet was then used for DNA extraction with the PROMEGA© Reliaprep gDNA kit and yielded DNA in the range of 5-80 ng µL^-1^. For species identification we used the 18S rRNA gene sequence (Primers 18S F 5′-ACCTGGTTGATCCTGCCAG-3′ and 18S R 5′-TGATCCTTCCGCAGGTTCAC-3′; 60°C, [26]) and PCR chemicals by PROMEGA© with the following PCR conditions: 2 min at 95 °C, 35 cycles with 30 s at 95 °C – 1 min at 60 °C – 30 s at 72 °C.

Viral genomic DNA was extracted by centrifuging 2 mL of lysate at maximum speed (20’000 rpm) for 10 minutes. The supernatant was discarded and DNA suspended in 30 µL Tris-EDTA buffer. We used primers for the DNA polymerase B gene (polB) (VpolAS4 GARGG[I]GC[I]AC[I]GT[I]YTNGA and VpolAAS1 CC[I]GTRAA[I]CCRTA[I]AC[I] SWRTTCAT) [27]. The same PCR chemicals as for the 18S rRNA gene was used but the PCR conditions were: 2 min at 95 °C, 35 cycles with 30 s at 95 °C – 1 min at 60 °C – 30 s at 72 °C.

All PCR products were cleaned up (Fisher Scientific PCR cleanup ®) prior to sequencing and the forward primer for each sequence was used for Sanger sequencing with EUROFINSGENOMICS®. The 18S rRNA gene sequences were clipped and trimmed for further analysis, and taxonomic identification was performed with Codon Code Aligner (Version 9.0.1. “CodonCode Aligner User Manual,” n.d.) and the following reference used: reference genome from [29], GenBank CAID01000001.2 to CAID01000020.2, RCC2590_scf16 BCC102-466838, Rcc809_18S_fromgenome, ostta12g00750, Olu_18S_fromgenome. The polB sequences were also clipped and trimmed for further analysis but we then used the NCBI Blast search tool for species identification. Both 18S rRNA and polB gene sequences were then used to build neighbour-joining phylogenetic trees (Codon Code Aligner) to identify relatedness. We used the sequencing results to compare to the infection patterns.

In addition to the taxonomic identity, we characterized both host and virus strains by their isolation origin, namely the origin of the water sample they were isolated from (geography and season). In addition, since viruses have an inherent host dependency, they were also characterized in relation to the water sample of their hosts. These characteristics will then be used to analyse the infection patterns in the following experiment.

### Cross infection experiments

In total, 17 *Ostreococcus* sp. strains (2 *O. tauri* and 15 *O. mediterraneus*) were infected by all 79 different *Ostreococcus* viruses. Specifically, we prepared duplicate (i.e. 2 drops of lysate per virus) cross infection experiments via plaque assays combining all *Ostreococcus* strains with all virus strains resulting in 1343 combinations. The multiplicity of infection (MOI) of all infections was 10:1 where the *Ostreococcus* cultures had a starting concentration of 5*10^6^ cells mL^-1^ in 0.2 % agarose (Sigma Aldrich ®) and 2 µL of virus lysate containing 50*10^6^ viral particles mL^-1^ were added (see Fig. S1 for petri dish schematic). Determination of abundance of viral particles was done via flow cytometry. The petri dishes were kept at 18 °C and a 12:12 day and night cycle.

After 3, 5, 7 and 10 days of incubation, we recorded plaque formation (presence/absence) and took photographs of the petri dishes using a Canon digital camera. Plaque size was measured via ImageJ ® (V 1.53t) [30] and the rate of plaque size increase over the 10 days post infection was calculated based on the slope of a linear regression fitted to the size data from day 3 to day 10 post infection.

### Statistical analysis

We used the bipartite network analysis to test for nestedness in the infection patterns using ‘computeMod()’ in the bipartite package in the R programming environment [31]. To test the significance of the patterns we calculated the nestedness metric based on overlap and decreasing fill (NODF) of 1000 null models and compared their distribution with the NODF of the observed matrix.

Due to non-normalized data in the rate of plaque size increase and unequal sample sizes we used non-parametric approaches (Wilcoxon-Mann-Whitney-Test) and subsequent post hoc tests for comparison between groups (between and within clusters) with Bonferroni corrections. All distance matrix calculations as well as statistical analyses were carried out using R software (R Version 4.2.2. Cran Server) and the following specific packages in the R environment: ggplot2, gridExtra, ggpubr, reshape2, nlme, rnaturalearth, bipartite, ggtree.

## Results

### Isolation success

Out of 119 water samples, we successfully isolated two and 21 strains of *Ostreococcus tauri* and *Ostreococcus mediterraneus*, respectively. These strains originated largely from the Kiel Bight and Mecklenburg Basin with a higher isolation success during spring and autumn phytoplankton blooms. The isolation success was approximately 7 % as we isolated *Ostreococcus* strains from 8 out of 119 water samples. Then, we were able to isolate 9 *Ostreococcus tauri* viruses (OtVs) and 70 *Ostreococcus mediterraneus* viruses (OmVs) from 13 of the *Ostreococcus* strains using 84 different water samples (Fig. S3 overview of virus isolations). The viruses mainly originated from the Kiel Bight (KB) and Mecklenburg Bight (MB) in spring and early summer (March 2018 to May 2020).

The 18S rRNA gene analysis (based on nucleotide sequence) identified the host strains to be *O. tauri* and *O. mediterraneaus* with single nucleotide genetic differences described already in [32] (Fig. 3a). The two *O. tauri* strains are grouped in the lower branch of the tree whereas the *O. mediterraneus* strains are grouped in the upper branch of the tree. The polB nucleotide sequence analysis of the viruses with NCBI Blast revealed three different virus species (Fig. 3b, *O. mediterraneus* virus (OmV), *O. tauri* virus (OtV) and *O. lucimarinus* virus (OlucV) in names). Note that even though the host of OlucV virus was *O. mediterraneus*, the polB sequence was most similar to that of a *O. lucimarinus* virus. Compared to the 18S rRNA gene sequences, we found more nucleotide variability among the viruses forming five distinct strain clusters (Fig. 3b).

**Figure 2.**
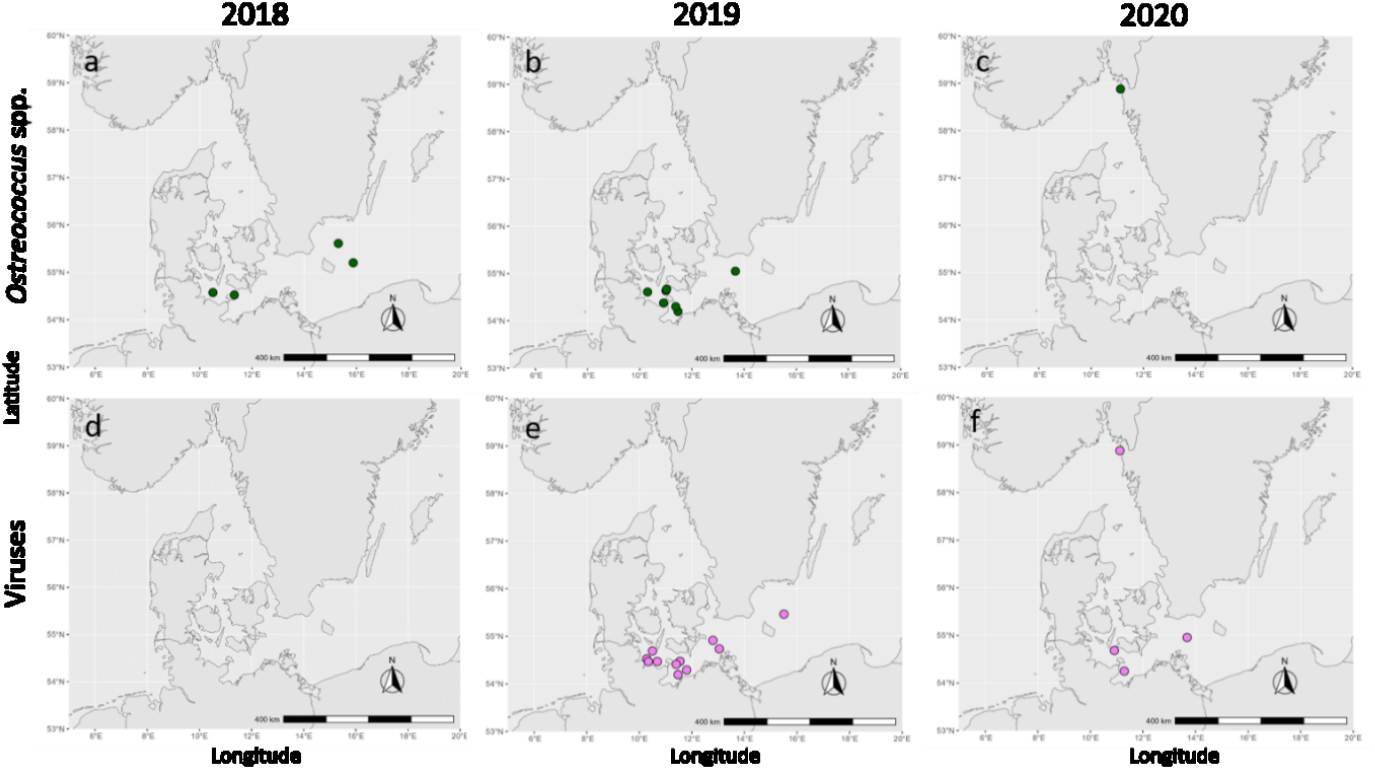
The panels show all stations from which we successfully isolated *Ostreococcus* sp. (green points in panels a-c) and their viruses (violet points in panels d-f). Note that we used *Ostreococcus* hosts in combination with all water samples from the same year to start the isolation procedure of viruses.

**Figure 3.**
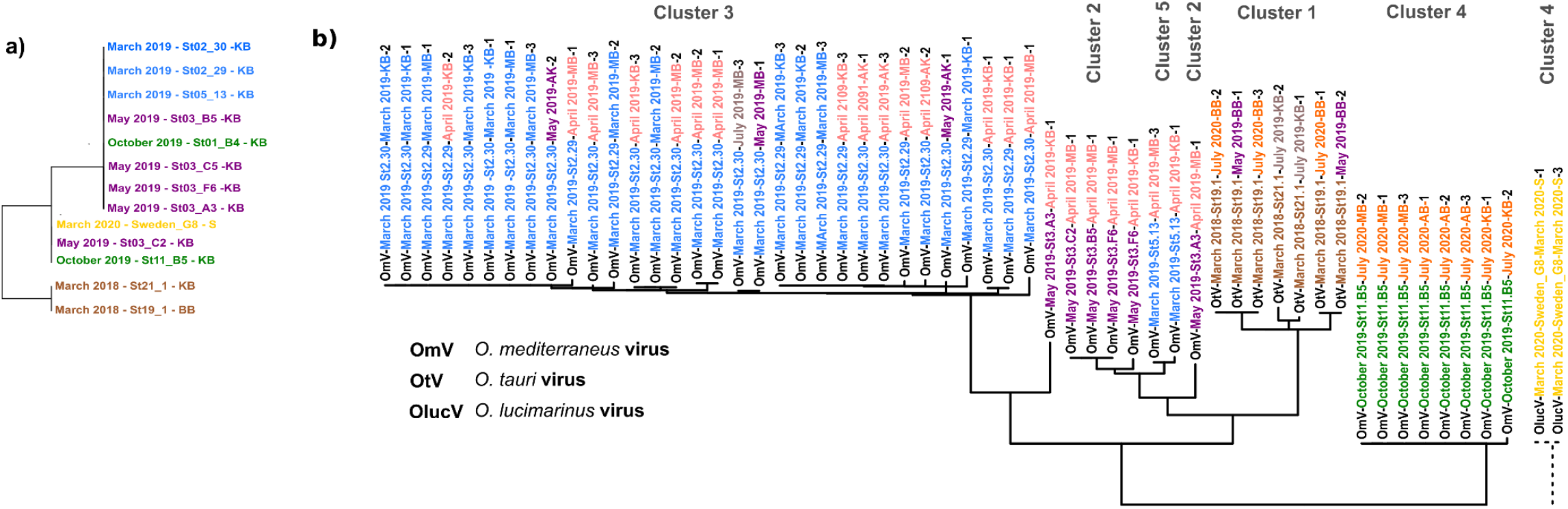
Panel a and b show the genetic trees of the *Ostreococcus* and virus strains based on 18S rRNA gene and DNA polymerase beta (PolB) sequences, respectively. Each of the virus strain names indicates the virus species, the isolation host, the seasonal and geographic origin of the water sample and the number of the viral strain of that combination (e.g. OmV-May 2019_St.3_A3-March 2019-MB-2) The colors indicate the different seasons from when the water samples were taken. Two- and one letter abbreviations at the end of the names indicate the geographical region where the water samples for virus isolations were taken (KB=Kiel Bight, MB=Meckelnburg Bight, AB=Arkona Basin, BB=Bornholm Basin, S=Sweden). The sequences of OlucV were too different to be aligned with the rest of the PolB sequences.

### Cross infection experiment

Of the 1343 host-virus combinations, we found successful infections in 206 combinations (Fig. 4a, blue squares in the infection matrix). These successful infections clustered into five modules (cluster 1-5) based on the modularity analysis in which we found different patterns in relation to both the hosts and viruses (Fig. 3b). In clusters 2 and 3, we found that *Ostreococcus* viruses that were isolated from hosts from a specific time of year, for example from March or May 2019, were generally able to infect other hosts from that same time period as well. Hosts from any other month were not infected. In clusters 1, 4, and 5, we found that the viruses were mainly host specific with the exception of some viruses (virus of host 21.1 and virus of host Sweden G8) being able to also infect hosts that were regionally far away. While the infection patterns are variable, we find that overall, we have a restriction of infections to species specificity and/or seasonal specificity. It is important to indicate that the modularity patterns relate to the viruses’ host origin but the virus water samples came from different times. For example: the difference between clusters 2 and 1 is that the virus water samples in cluster 2 were from the same season (May 2019 vs. April 2019) whereas in cluster 1 the virus water samples were from one or two years later (March 2018 vs. May, July 2019 and July 2020). However, these virus seasonal origins did not affect the modularity pattern.

**Figure 4.**
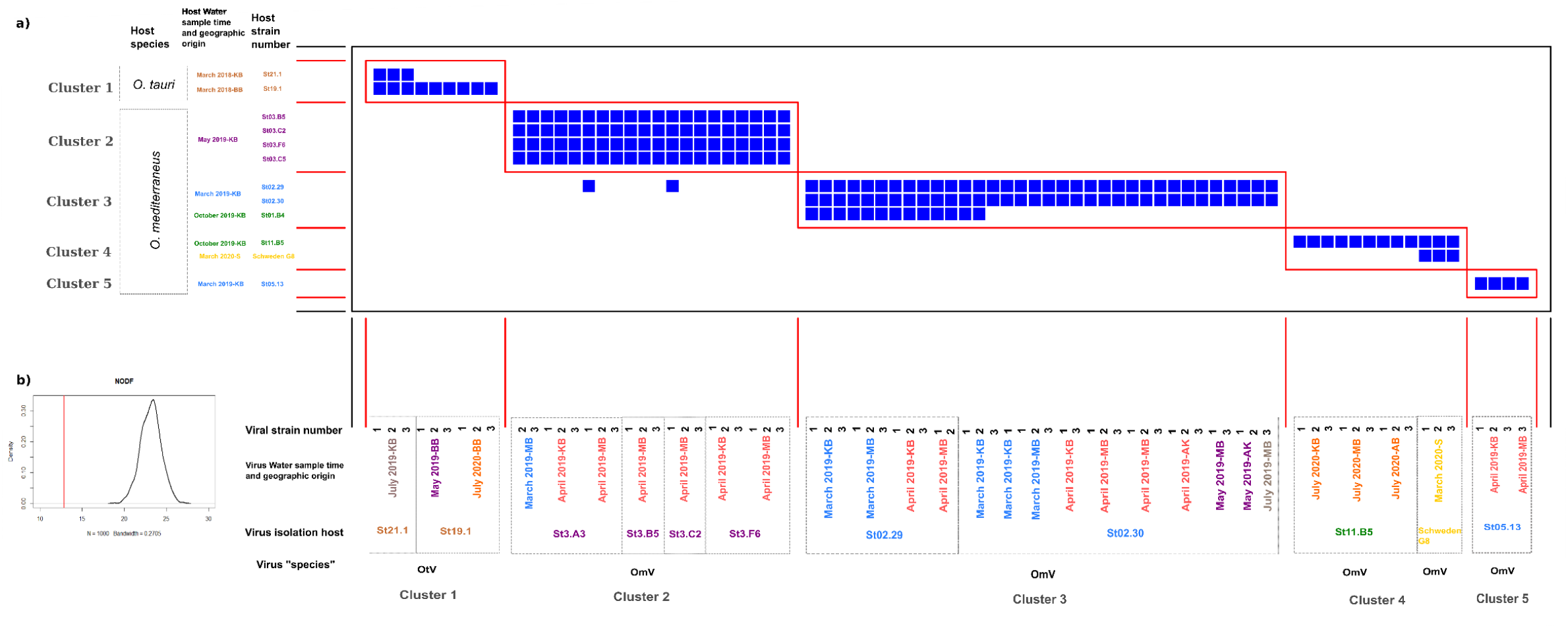
Panel a shows the infection matrix after the modularity analysis (note that only strains where infection was successful are shown in the matrix). The virus strains are shown on the x-axis whereas the host strains are shown on the y-axis. The colours on the axes’ labels indicate the timing of the cruises from which the hosts were isolated or from which the water samples of the viruses were collected. Blue squares framed with red lines indicate successful infection clusters. Panel b shows the results of the NODF distribution of the calculated null models and the observed NODF (red line) indicating that the depicted modularity of the infection matrix was significant.

Within the different clusters some plaques appeared on day 3 post infection, whereas others only appeared on the 5^th^ or 7^th^ day post infection (Fig. S4). At the same time, the plaques grew with different velocities (Fig. S5 slope of regression lines). Of the successful infections, we calculated the rate of increase in plaque size per day and checked for differences between the clusters and the difference of isolation-host infections and foreign host infections. We found significant differences between the clusters (Chi^2^= Clusters: _4_ = 41.506, p < 0.001). Specifically, we find a significant difference for the host species being infected in Cluster 3 and 5 (Fig. 5c Wilcoxon Tests, p=0.0045 and p=0.1, respectively). Overall from the first to last cluster the rate of plaque size change decreases by about 50%. In addition, we found that the variability within the virus-host infectivity clusters was highest in cluster 2 (Fig. 5c). This variation between and within the clusters indicates ecological differences between the host-virus pairs and their related infection dynamics.

**Figure 5.**
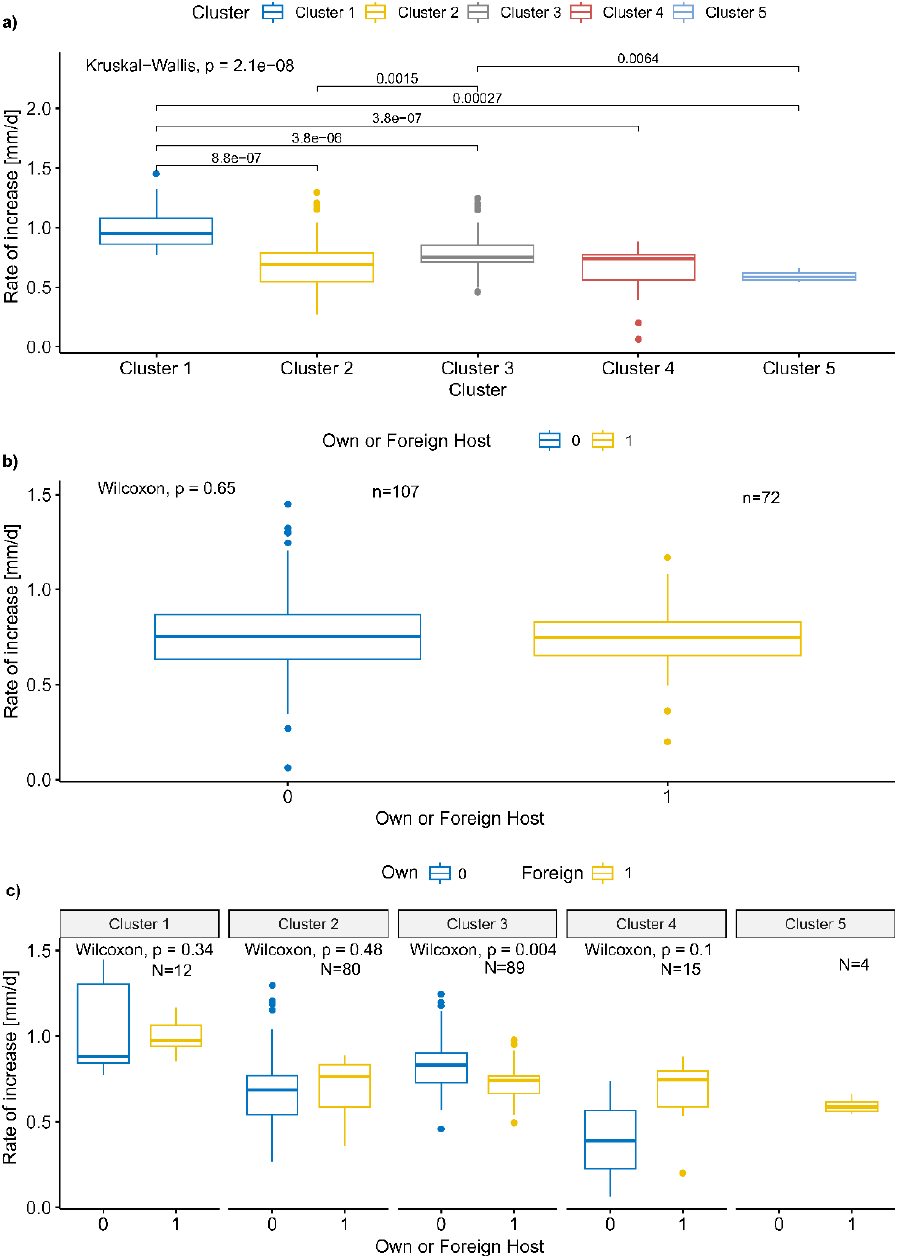
The figure shows the rate of increase plaque sizes of the successful infections for each of the clusters of virus-host infection separately in panel a, for the difference between own and foreign hosts in panel b and then divided into the clusters in panel c. In panel b and c the colors indicate original host and foreign hosts. The boxplots indicate the mean and 95% confidence interval. In each panel and subpanel the significance results of a non-parametric rank test is shown comparing the rate of increase between the hosts (panel b and c) and between the clusters (panel a).

## Discussion

Over a two-year sampling period, we were able to isolate 23 hosts and 79 virus strains of the *Ostreococcus* sp. species complex from different regions of the Baltic Sea as well as the Swedish Skagerrak and investigated infection patterns within this system. We identified the evolutionary history, and thus the timing of when hosts and their associated viruses coexisted, as the main driver of infection patterns. In addition, host species and strain specificity underline the present understanding of rapid host-virus co-evolution [33, 34], and here, we demonstrate this in a phytoplankton-virus system that evolved in natural conditions.

Based on metagenomic analyses, *Ostreococcus* sp. is present all year round in the Baltic Sea, but its presence varies throughout the year and regionally [35]. This is in line with the isolation success we had over the two-year sampling period where we were mainly able to isolate *Ostreococcus* strains in spring (March to May) and autumn (October). Spring blooms in the Western Baltic Sea are dominated by large and often chain forming diatoms [20] that can be easily separated via sieving from the smaller picoplankton fraction. When applying nutrient enrichment excluding silicate and growth at lower (ca. 15 °C) temperatures, *Ostreococcus* sp. is favoured over other picoplankton species and diatoms for isolation approaches. In contrast, during summer blooms the Western Baltic Sea phytoplankton community is dominated by cyanobacteria [36]. These cyanobacteria are the dominant picoplankton species in summer and hinder the isolation success of *Ostreococcus* sp. in this season. Our findings on timely biases for isolation success have also been observed in the Mediterranean *Ostreococcus* populations that are mainly present in early times of the year in the Mediterranean Sea. Despite the molecular evidence for the presence of genetic material of *Ostreococcus* even in the Eastern part of the Baltic Sea [35], our success of isolating *Ostreococcus* was confined to the Western Baltic. As *Ostreococcus* are primarily marine species, the pronounced brackish conditions in the Eastern Baltic Sea could hinder prevalence of algae in that region [37]. In summary, for the successful isolation of phytoplankton species knowing when the species of interest is likely to be either dominant in the community or easily separable from other community members is crucial.

The 18S rRNA gene sequences revealed that our *Ostreococcus* species belonged mainly to the subclade D, also named *O. mediterraneus*, while two strains belonged to the *O. tauri* clade C [17, 32]. This is surprising because for example Zeigler et al. (2017) found that most of the *Ostreococcus* sequences were identified as *O. tauri*. Within the species complex of *Ostreococcus*, there are very few base pair differences within the 18S rRNA gene [32], and since *O. tauri* was the first species to be isolated and identified within this picoplankton species complex [38], available genomic data could actually underestimate the prevalence of each of the four subclades A-D [39].

The genetic relatedness of the isolated virus strains based on Pol B sequences (typically used gene for identification and phylogenetic classification [40]) showed a clustering into groups based on the host’s origin and the infection clusters. This is in line with a previous study showing that genetically related viruses tend to infect the same hosts or group of hosts [11]. The pol B sequence of the two viral strains originating from the Sweden_G8 *O. mediterraneus* host, did not align with the rest of the sequences because they were more similar to *O. lucimarinus* pol B sequences based on BLAST comparisons. This was a contradictory result [41] because following the “identification” of the virus based on its host origin (ecological identification), we would have had to call the virus an *O. mediterraneus* virus. This indicates that the genetic identification of viruses may not be adequate to deduce its host origin or its role in the “ecosystem”. We will now further discuss the ecology of the infections in this study considering the viruses host origin.

In previous studies, different virus or host characteristics have been described ranging from generalist to specialist viruses or highly susceptible to highly resistant hosts. The thus emergent infection patterns based on the virus or host characteristics have been described in [42]. Clusters of infection form when a set of hosts or viruses exhibit the same traits with regards to infectability or resistance. Identifying what drives modularity in infection networks is key to understanding the ecology of the infection dynamics in a system. In our study, we found that the modules in the infection patterns were not random (see Fig. 3, panel b). Some viral clusters could be associated to the time of when hosts were isolated and some infections were strain-specific but all except one were species-specific. Specifically, for the clusters associated to seasons we found that the viruses that were isolated from hosts from a specific time of year (March or May) were also able to infect hosts from that same time of year. However, all other hosts that were tested, were not infected. The strain/species specificity underlines previous findings on host specificity in the *Ostreococcus* virus system [11]. The seasonal modularity however, indicates co-evolutionary patterns in this host-virus system on a time scale that has previously been described and studied in another phytoplankton-virus system [11, 33, 34].

While it is known that viruses are species-specific, infections could also be specific to environmental conditions [43], and we could therefore find that viruses are specifically able to infect only hosts from similar seasons when for example temperature conditions stay within a specific range (see for example clusters 3 and 4 and see Fig. S6 for changes in abiotic environment over time). In addition to regular saline water inflow from the North Sea, the Baltic Sea experiences an increasing inflow of water from coastal freshwater sources (related to precipitation and in spring after snow melt) inducing stratification [44], which in turn leads to increasing temperature in upper water layers. These conditions might also allow *Ostreococcus* spp. to flourish [45]. Being able to cross-infect in spring/March and independent of changes in salinities could be an important asset for viruses of *Ostreococcus* to take advantage of thriving subpopulations of the host.

*Ostreococcus* spp. are non–blooming picoplankton species that are adapted to the environmental conditions (such as temperature, salinity and nutrients) that they have experienced in the past (i.e. several 10s to hundreds of generations, see Listmann et al. (2020) for the example of carbon uptake). These adaptations do not only relate to physiological or fitness related traits but likely also the biotic interactions with viruses. Between the seasons from which the hosts were isolated and which related to the infection modules the temperature, sometimes salinity but also light conditions varied (see supplement of Zhong et al. 2020) (Santelia et al 2022 preprint). For instance, day-to-day salinity variations in the upper few centimetres of the water column during the sampling campaign in the Swedish Skagerrak ranged from 22.5 to 30.0 (supplement of Rahlff et al. (2022). The virus OlucV_March 2020_Sweden_G8_March_2020_S3 isolated from a water sample from this station was still capable of infecting hosts from the Western Baltic Sea, which are usually subjected to much lower salinities (Fig S6 panel a vs panel b; Western vs. Eastern sampling area). Another example of viruses from a higher salinity environment that were able to infect a host from a lower salinity environment are OtV_March 2018_St21.1_July 2019_KB. These examples could reflect the viruses’ adaptation to salinity gradients, which seemingly did not negatively impact host range and infection capabilities. In line with that, a study by Bellec et al. (2010) showed that coastal marine lagoons, which due to their shallowness are also more prone to salinity changes, contained *O. tauri* viruses that were also abundant in offshore and coastal ecosystems.

Further infection experiments with changes in salinity or temperature as compared to pure co-evolutionary studies in this system would shed light on these two possible explanations of the infection patterns we found. In addition, finding differences in infection patterns depending on the environmental condition, could allow us to make predictions how phytoplankton would cope under future climate conditions.

The fact that we find the plaque size increase to vary even within some clusters of infection shows that though we have clusters of infection, viruses could still be specifically adapted to their hosts and the strength of infection is not always the same. We are not aware of studies in (pico)-phytoplankton that have specifically investigated strengths of infection in a temporal scale in such a large infection assay (except maybe Bellec et al. 2014). The differences in plaque size change and when the plaques appear could be used to identify several aspects of the infections: i) the later a plaque becomes visible, the potentially longer the latency period from the start of infections is. ii) the faster a plaque increases, the faster the virus spreads and iii) the larger the plaque becomes, the stronger the infection is. However, liquid infection assays where host and virus traits are usually identified [48] are needed to validate these assumptions. All of the mentioned characteristics of the infection will affect the infection dynamics within the ecosystem [10] and potential effects on the *Ostreococcus* populations [49]. Therefore future studies should consider incorporating and experimenting on the documentation of these infection attributes to better characterize and identify infection patterns related to the seasonal and geographical origin of both the hosts and viruses.

## Supporting information

Supplementary Material Listmann et al

## Acknowledgements

We like to thank Stefanie Schnell, Tabea Schneider and Gwenael Pigneau for contributions to the experimental work as well as scientific input. We like to thank the Sven Lovén Centre for Marine Sciences, Tjärnö, Sweden as part of the University of Gothenburg for hosting JR and SPE during the MIDSEAS campaign and providing experimental and sampling facilities.

## References

1. Pomeroy LR, le Williams PJB, Azam F, Hobbie JE (2007) The microbial loop. Oceanography 20:28–33. https://doi.org/10.5670/OCEANOG.2007.45

2. Wilhelm SW, Suttle CA (1999) Viruses and Nutrient Cycles in the Sea aquatic food webs. Bioscience 49:781–788. https://doi.org/VirusesandNutrientCyclesintheSeaaquaticfoodwebs

3. Suttle CA (2007) Marine viruses - Major players in the global ecosystem. Nat Rev Microbiol 5:801–812. https://doi.org/10.1038/nrmicro1750

4. Coutinho FH, Gregoracci GB, Walter JM, et al (2018) Metagenomics Sheds Light on the Ecology of Marine Microbes and Their Viruses. Trends Microbiol. 26:955–965

5. Castillo YM, Sebastián M, Forn I, et al (2020) Visualization of Viral Infection Dynamics in a Unicellular Eukaryote and Quantification of Viral Production Using Virus Fluorescence in situ Hybridization. Front Microbiol 11:. https://doi.org/10.3389/fmicb.2020.01559

6. Gwak HJ, Rho M (2022) ViBE: a hierarchical BERT model to identify eukaryotic viruses using metagenome sequencing data. Brief Bioinform 23:. https://doi.org/10.1093/BIB/BBAC204

7. Kieft K, Adams A, Salamzade R, et al (2022) vRhyme enables binning of viral genomes from metagenomes. Nucleic Acids Res 50:e83–e83. https://doi.org/10.1093/NAR/GKAC341

8. Johannessen TV, Larsen A, Bratbak G, et al (2017) Seasonal Dynamics of Haptophytes and dsDNA Algal Viruses Suggest Complex Virus-Host Relationship. Viruses 2017, Vol 9, Page 84 9:84. https://doi.org/10.3390/V9040084

9. Yau S, Caravello G, Fonvieille N, et al (2018) Rapidity of genomic adaptations to prasinovirus infection in a marine microalga. Viruses 10:1–11. https://doi.org/doi:10.3390/v10080441

10. Correa AMS, Howard-Varona C, Coy SR, et al (2021) Revisiting the rules of life for viruses of microorganisms. Nat. Rev. Microbiol. 19:501–513

11. Clerissi C, Desdevises Y, Grimsley N (2012) Prasinoviruses of the Marine Green Alga Ostreococcus tauri Are Mainly Species Specific. J Virol 86:4611–4619. https://doi.org/10.1128/JVI.07221-11

12. Nagasaki K, Yamaguchi M (1997) Isolation of a virus infectious to the harmful bloom causing microalga Heterosigma akashiwo (Raphidophyceae). Aquat Microb Ecol Aquat Microb Ecol 13:135–140

13. Kranzler CF, Krause JW, Brzezinski MA, et al (2019) Silicon limitation facilitates virus infection and mortality of marine diatoms. Nat Microbiol 4:1790–1797. https://doi.org/10.1038/s41564-019-0502-x

14. Mojica KDA, Brussaard CPD (2014) Factors affecting virus dynamics and microbial host-virus interactions in marine environments. FEMS Microbiol Ecol 89:495–515

15. Demory D, Arsenieff L, Simon N, et al (2017) Temperature is a key factor in Micromonas-virus interactions. ISME J 11:601–612. https://doi.org/10.1038/ismej.2016.160

16. Harley CDG, Randall Hughes a, Hultgren KM, et al (2006) The impacts of climate change in coastal marine systems. Ecol Lett 9:228–41. https://doi.org/10.1111/j.1461-0248.2005.00871.x

17. Demir-Hilton E, Sudek S, Cuvelier ML, et al (2011) Global distribution patterns of distinct clades of the photosynthetic picoeukaryote Ostreococcus. ISME J 5:1095–1107

18. Bellec L, Grimsley N, Derelle E, et al (2010) Abundance, spatial distribution and genetic diversity of Ostreococcus tauri viruses in two different environments. Environ Microbiol Rep 2:313–321. https://doi.org/10.1111/j.1758-2229.2010.00138.x

19. Derelle E, Monier A, Cooke R, et al (2015) Diversity of Viruses Infecting the Green Microalga Ostreococcus lucimarinus. J Virol 89:5812–5821. https://doi.org/https://doi.org/10.1128/JVI.00246-15

20. Snoeijs-Leijonmalm P, Schubert H, Radziejewska T (2017) Biological oceanography of the Baltic Sea

21. Reusch TBH, Dierking J, Andersson HC, et al (2018) The Baltic Sea as a time machine for the future coastal ocean. Sci Adv 4:. https://doi.org/10.1126/sciadv.aar8195

22. Listmann L, Kerl F, Martens N, Schaum C-E (2020) Carbon acquisition in a Baltic pico-phytoplankton species-Where does the carbon for 1 growth come from? 2. bioRxiv 2020.09.07.285478. https://doi.org/10.1101/2020.09.07.285478

23. Rahlff J, Esser SP, Plewka J, et al (2022) Heads in the clouds : marine viruses disperse bidirectionally along the natural water cycle. bioRxiv

24. Anderson RA, Berges RA, Harrison PJ, Watanabe MM (2005) Recipes for Freshwater and Seawater Media; Enriched Natural Seawater Media. Algal Cult Tech 596

25. (2017) The Nature of Viruses. Fenner’s Vet Virol 3–16. https://doi.org/10.1016/B978-0-12-800946-8.00001-5

26. Cai H, Wang K, Huang S, et al (2010) Distinct patterns of picocyanobacterial communities in winter and summer in the chesapeake bay. Appl Environ Microbiol 76:2955–2960. https://doi.org/10.1128/AEM.02868-09/ASSET/65CD8D44-968B-4554-B641-62072C5D62D7/ASSETS/GRAPHIC/ZAM9991009100003.JPEG

27. Clerissi C, Grimsley N, Ogata H, et al (2014) Unveiling of the diversity of prasinoviruses (Phycodnaviridae) in marine samples by using high-throughput sequencing analyses of PCR-amplified DNA polymerase and major capsid protein genes. Appl Environ Microbiol 80:3150–3160. https://doi.org/10.1128/AEM.00123-14

28. CodonCode Aligner User Manual

29. Derelle E, Yau S, Moreau H, Grimsley NH (2018) Prasinovirus Attack of Ostreococcus Is Furtive by Day but Savage by Night. J Virol 92:e01703–17. https://doi.org/10.1128/JVI.01703-17

30. Schneider CA, Rasband WS, Eliceiri KW (2012) NIH Image to ImageJ: 25 years of image analysis. Nat Methods 2012 97 9:671–675. https://doi.org/10.1038/nmeth.2089

31. Carsten F. Dormann bipartite-package: Analysis of bipartite ecological webs in bipartite: Visualising Bipartite Networks and Calculating Some (Ecological) Indices. https://rdrr.io/cran/bipartite/man/bipartite-package.html. Accessed 10 Nov 2022

32. Subirana L, Péquin B, Michely S, et al (2013) Morphology, genome plasticity, and phylogeny in the genus ostreococcus reveal a cryptic species, o. mediterraneus sp. nov. (mamiellales, mamiellophyceae). Protist 164:643–659. https://doi.org/10.1016/j.protis.2013.06.002

33. Frickel J, Sieber M, Becks L (2016) Eco-evolutionary dynamics in a coevolving host-virus system. Ecol Lett 19:450–459. https://doi.org/10.1111/ele.12580

34. Koch H, Frickel J, Valiadi M, Becks L (2014) Why rapid, adaptive evolution matters for community dynamics. Front Ecol Evol 2:1–10. https://doi.org/10.3389/fevo.2014.00017

35. Zeigler Allen L, McCrow JP, Ininbergs K, et al (2017) The Baltic Sea Virome: Diversity and Transcriptional Activity of DNA and RNA Viruses. mSystems 2:e00125–16. https://doi.org/10.1128/mSystems.00125-16

36. Eilola K, Meier HEM, Almroth E (2009) On the dynamics of oxygen, phosphorus and cyanobacteria in the Baltic Sea; A model study. J Mar Syst 75:163–184. https://doi.org/10.1016/j.jmarsys.2008.08.009

37. Clayton S, Lin YC, Follows MJ, Worden AZ (2017) Co-existence of distinct Ostreococcus ecotypes at an oceanic front. Limnol Oceanogr 62:75–88. https://doi.org/10.1002/LNO.10373

38. Courties C, Vaquer A, Troussellier M, et al (1994) Smallest eukaryotic organism. Nature 370:255–255. https://doi.org/10.1038/370255a0

39. Tragin M, Vaulot D (2019) Novel diversity within marine Mamiellophyceae (Chlorophyta) unveiled by metabarcoding. Sci Rep 9:1–14. https://doi.org/10.1038/s41598-019-41680-6

40. Cassedy A, Parle-McDermott A, O’Kennedy R (2021) Virus Detection: A Review of the Current and Emerging Molecular and Immunological Methods. Front Mol Biosci 8:76. https://doi.org/10.3389/FMOLB.2021.637559/BIBTEX

41. Van Regenmortel MHV (2007) Virus species and virus identification: Past and current controversies. Infect Genet Evol 7:133–144. https://doi.org/10.1016/j.meegid.2006.04.002

42. Flores CO, Meyer JR, Valverde S, et al (2011) Statistical structure of host-phage interactions. Proc Natl Acad Sci U S A 108:. https://doi.org/10.1073/pnas.1101595108

43. Maat DS, Biggs T, Evans C, et al (2017) Characterization and temperature dependence of arctic micromonas polaris viruses. Viruses 9:6–9. https://doi.org/10.3390/v9060134

44. Lass HU, Matthäus W (2008) General Oceanography of the Baltic Sea. State Evol Balt Sea, 1952-2005 A Detail 50-Year Surv Meteorol Clim Physics, Chem Biol Mar Environ 5–43. https://doi.org/10.1002/9780470283134.CH2

45. Guyon JB, Vergé V, Schatt P, et al (2018) Comparative Analysis of Culture Conditions for the Optimization of Carotenoid Production in Several Strains of the Picoeukaryote Ostreococcus. Mar Drugs 16:. https://doi.org/10.3390/MD16030076

46. Zhong D, Listmann L, Santelia M-E, Schaum C-E (2020) Functional redundancy in natural pico-phytoplankton communities depends on temperature and biogeography. Biol Lett 16:20200330. https://doi.org/10.1098/rsbl.2020.0330

47. Bellec L, Clerissi C, Edern R, et al (2014) Cophylogenetic interactions between marine viruses and eukaryotic picophytoplankton. BMC Evol Biol 14:1–13. https://doi.org/10.1186/1471-2148-14-59

48. Lievens EJP, Agarkova I, Dunigan DD, et al (2022) Life history diversity and signals of trade-offs in a large group of chloroviruses

49. Talmy D, Beckett SJ, Taniguchi DAA, et al (2019) An empirical model of carbon flow through marine viruses and microzooplankton grazers. Environ Microbiol 21:2171–2181. https://doi.org/10.1111/1462-2920.14626

