## Supplementary Material Listmann et al for "Co-evolutionary patterns shown in *Ostreococcus*-virus system from the Western Baltic Sea in freshly isolated hosts and viruses"

Supplementary material to Manuscript Listmann et al.

*Flow Cytometry*

*Ostreococcus* cells were counted at a rate of 65 µl $min^{-1}$ on a flow cytometer (Accuri BD C6 Plus) using gating of cells characterized by the height of forward scatter and red fluorescence signal (Fig. S2). The forward scatter was set to a threshold of ca. 200 that allowed us to determine debris and potentially high amounts of bacteria in the experimental cultures. *Ostreococcus* is large enough to be well distinguished from bacterial cells or cell debris (Fig. S2). To count viral particles via flow cytometry we adapted the protocol by Brussaard, Corina, (2004) the following way: virus samples (diluted or undiluted) were combined with Glutaraldehyde to a final concentration of 1% and after 15 min of incubation at room temperature, frozen at -80 °C for later analysis. After quick defrosting at 37 °C, 40 µL of virus sample was combined with 960µl of a mix of 0.02 µM filtered (Whatman Anotop ®) Tris-EDTA (Sigma Aldrich ®) buffer and SybrGold (Thermofisher ®) dye (2.5 µL 1´000x SybrGold per 1 mL TE buffer). The mixture was then heated to 80 °C for 10 min and cooled at room temperature for another 15 min. After cooling, virus particles were counted at a rate of 17 µL $min^{-1}$ in the flow cytometer. The FITC channel for green fluorescence was set to 20 and the virus populations identified using the side scatter and FIT-C signal (Fig. S2).

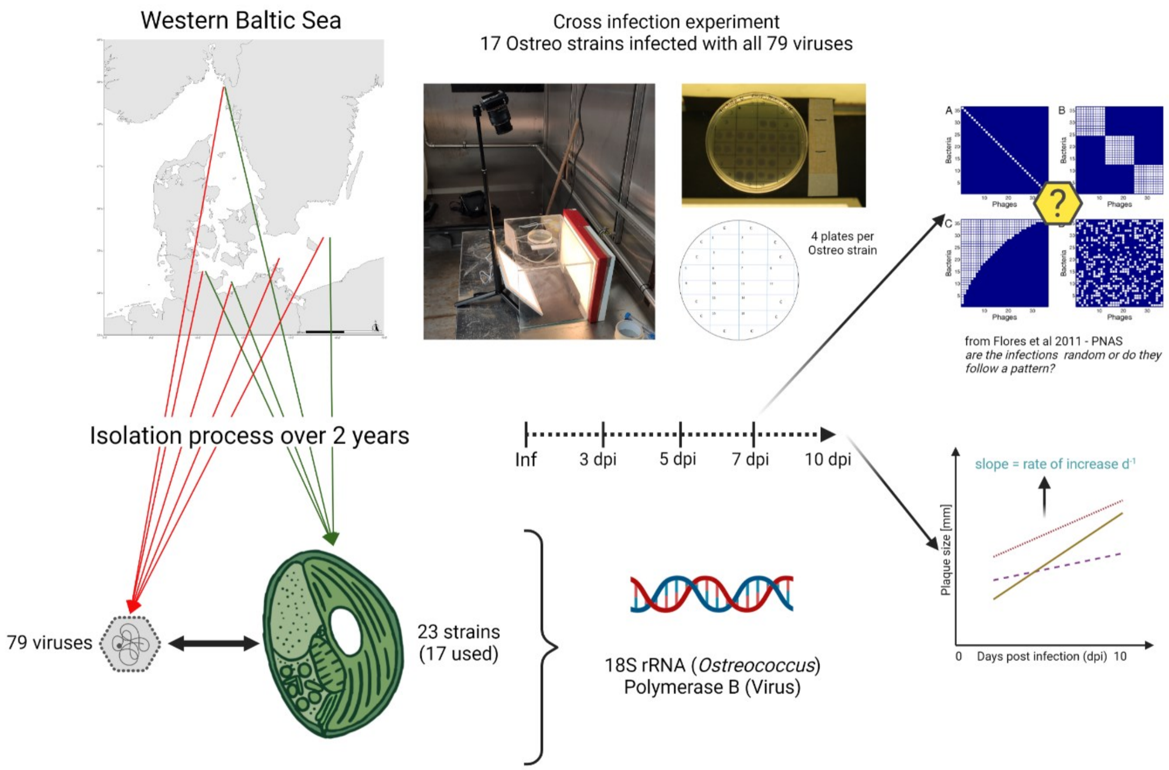


**Figure S1.** The isolation procedure and set up of the cross infection analysis as well as the genetic identification of both hosts and viruses is shown here. The modularity analysis was done 7 days post infection (dpi), i.e. the latest time point when infections appeared, whereas the calculation of the plaque size changes was done over 4 time points post infection (3, 5, 7 and 10 dpi). Figure was created using BioRender web application

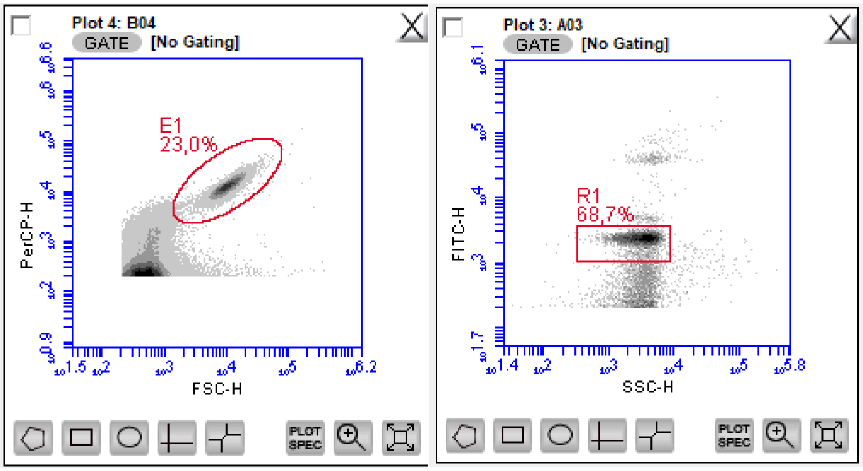


**Figure S2**: Flow Cytometry plot showing *Ostreoccocus* population (left panel) and virus populations (right panel). Left plot: the x-axis depicts the forward scatter which is a proxy for size and the y-axis depicts the blue fluorescence which is an indicator for chlorophyll a content. Right plot: the x-axis depicts the side scatter which is a measure for granularity and can also be used as a proxy for size and the y-axis depicts the green fluorescence for anything dyed with SybrGold.

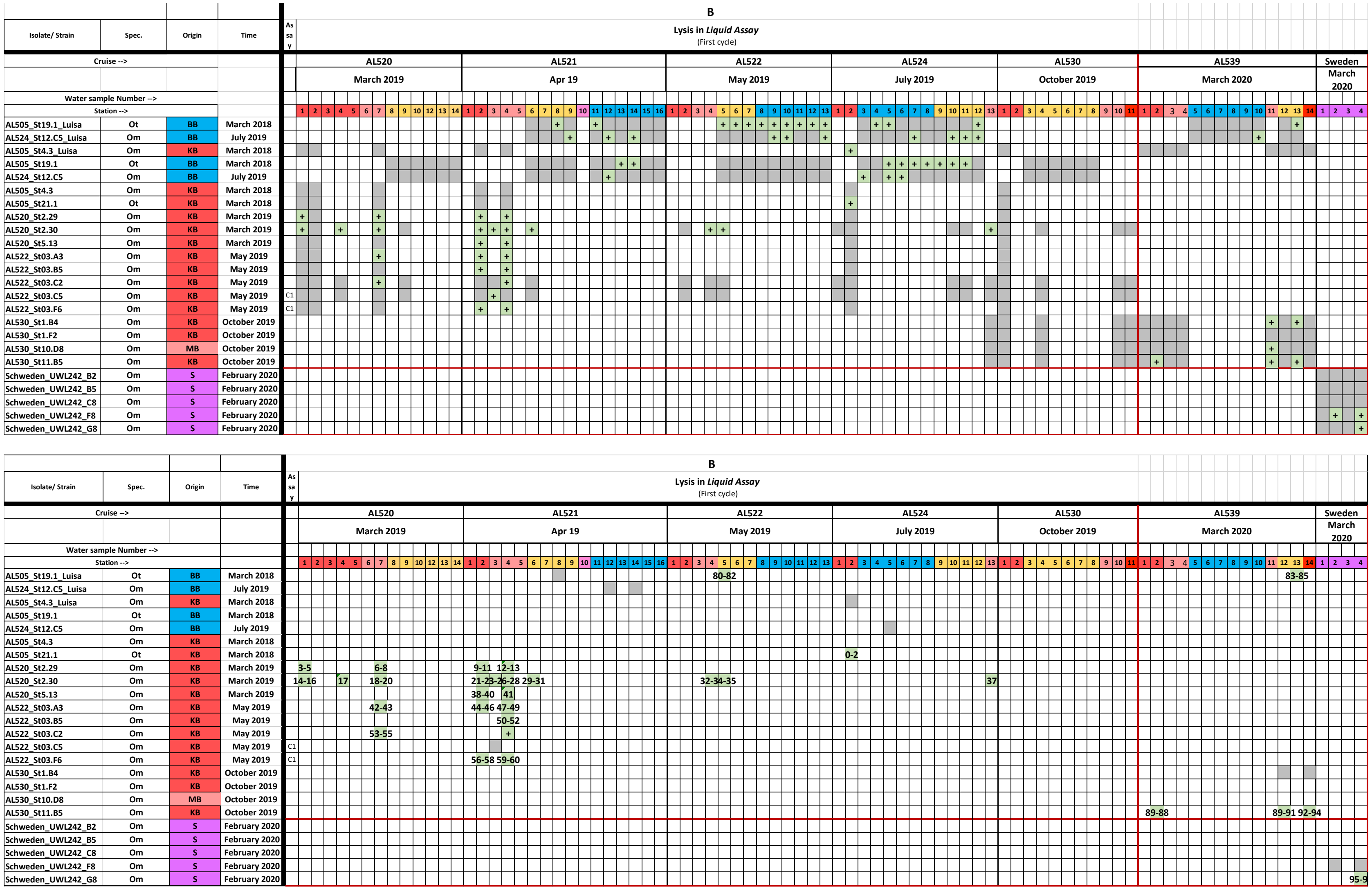


**Figure S3.** Top panel shows all *Ostreococcus* strains (left side) from which we started the isolation process for viruses. Grey boxes indicate a combination of *Ostreococcus* strain with water sample whereas green boxes indicate a successful lysis. Lower panel shows all remaining water and *Ostreococcus* combinations from which virus lysates remained and were used for the experiments.

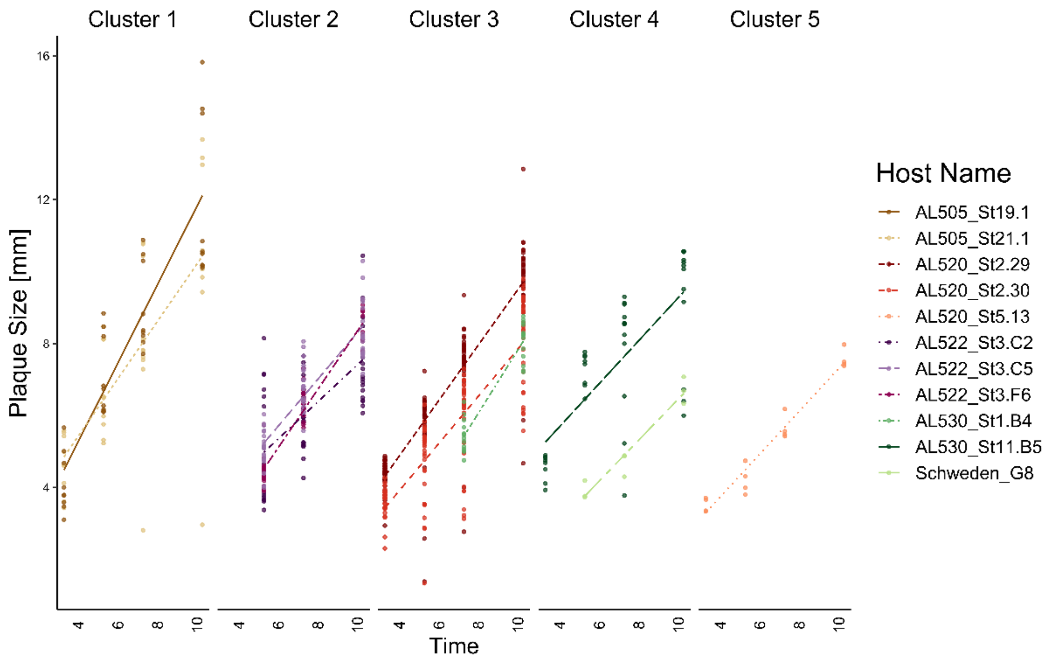


**Figure S4.** Points represent the plaque sizes in mm of the infections. Lines represent the linear regression fits for each host strain. The linetypes refer to the different hosts. The slopes indicate the rates of plaque size increase. Each panel shows the groups of infections belonging to one of the clusters in the infection matrix (infections per cluster range between 4 and 80 single infections.

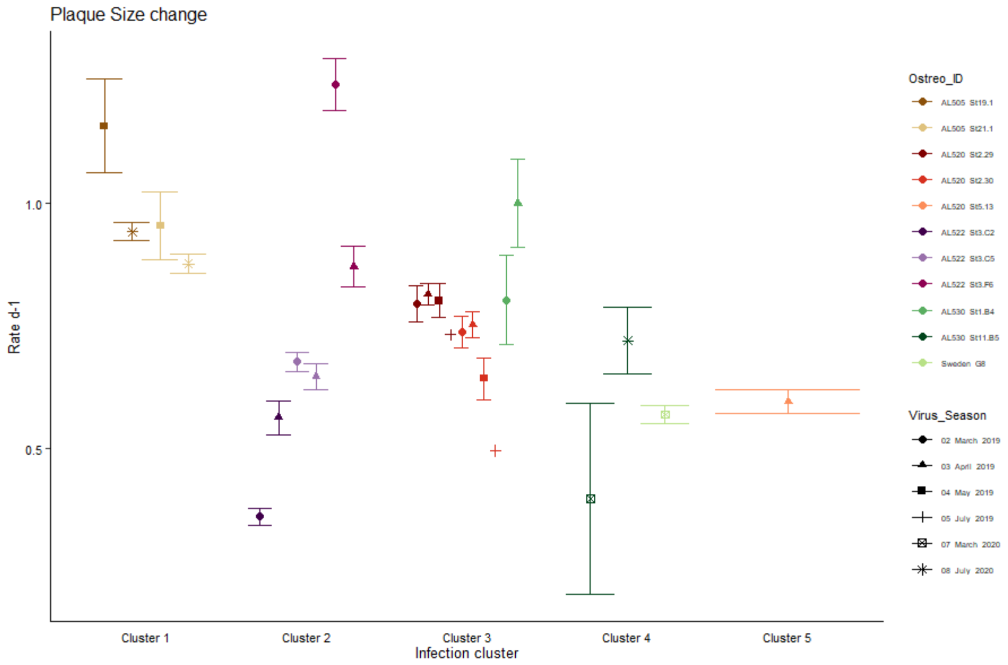


**Figure S5.** The figure shows the rate of change for the plaque sizes of the successful infections for each of the clusters of infection separately. The colors indicate the different host strains and the hues between the colors show the different strains isolated within one season. The different shapes indicate the season when viruses got extracted. The points and error bars show mean +/- SE.

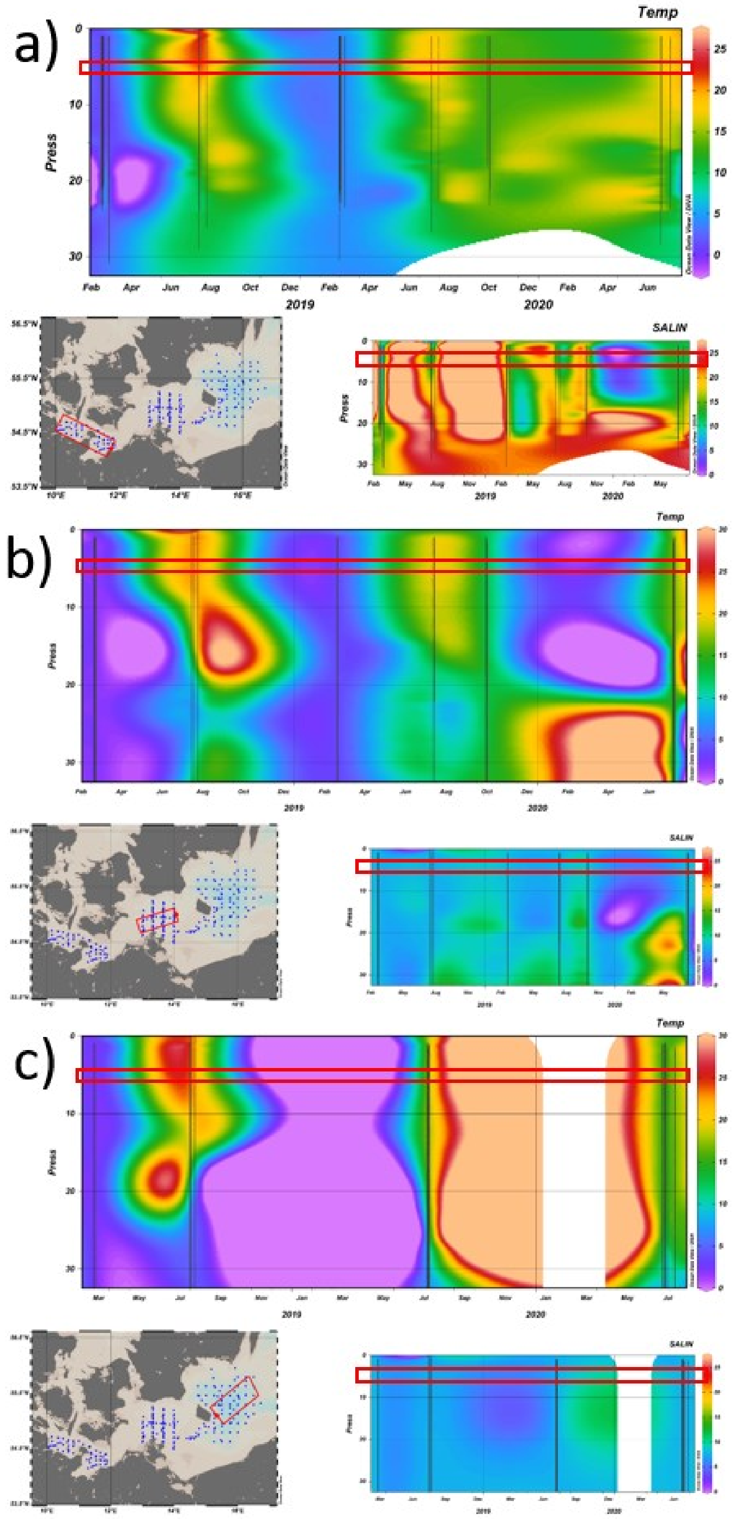


**Figure S6.** The figure shows the changes in temperature (bigger plot) and salinity (smaller plot) during the time of the sampling cruises. The different panels show the different research areas: Kiel Bight and Mecklenburg Basin KB and MB (a), Arkona Basin AB (b) and Bornholm Basin BB (c). The y-axes of the plots indicate the depth of the water, x-axes show the time of year during which the cruises took place and the colors indicate temperature °C and salinity PSU, respectively. In all basins temperature changes throughout the year and according to seasons, whereas salinity is rather constant over time and differs between the different geographical regions.
